# Computational discovery of precision therapeutics for hidradenitis suppurativa

**DOI:** 10.64898/2026.02.23.706050

**Authors:** Ernest Y. Lee, Phoebe Leboit, Haley B. Naik, Alice S. Tang, Francesco Vallania, Ashley E. Yates, Daniel M. Klufas, Scott L. Hansen, Michael D. Rosenblum, Margaret M. Lowe, Marina Sirota

## Abstract

Hidradenitis suppurativa (HS) is an underdiagnosed chronic, immune-mediated inflammatory skin disease that causes severe pain, drainage, and scarring, leading to significant physical and psychosocial burdens. HS is characterized by heterogenous molecular changes that are poorly understood, posing a significant challenge for drug development. Therapeutic options remain limited, and many patients experience disease relapse despite treatment. Therefore, precision medicine approaches are urgently needed to identify new therapies for HS. Here, we combine integrative transcriptomics, large-scale drug perturbational datasets, and translational immunology to identify sirolimus, pioglitazone, and fulvestrant as novel therapies for HS that can directly target and reverse the HS disease gene signature in immune cell types relevant to HS pathogenesis. Using a novel *ex vivo* HS skin model, sirolimus, pioglitazone, and fulvestrant inhibited T cell proliferation and activation, and suppressed the production of pro-inflammatory cytokines from HS skin. These results show that unbiased data-driven precision medicine approaches can identify novel therapies for HS and can serve more generally as a model approach for therapeutic discovery in other chronic inflammatory diseases.

**One Sentence Summary:** Data-driven precision medicine approach identifies sirolimus, pioglitazone, and fulvestrant as novel therapies for hidradenitis suppurativa

## Introduction

Hidradenitis suppurativa (HS) is a complex immune-mediated systemic inflammatory condition for which precision medicine is greatly needed. HS is characterized by chronic recurrent and painful inflammatory nodules and abscesses in intertriginous areas that can lead to irreversible fibronodular scarring and tunnel development, substantially impacting quality of life and mental health (*1*). HS is often associated with other chronic inflammatory skin conditions, such acne conglobata, dissecting cellulitis of the scalp, and pilonidal sinus, and cardiovascular comorbidities such as obesity, atherosclerosis, and type 2 diabetes (*2, 3*). The median diagnostic delay for HS is 7-10 years, and many patients present with late-stage disease due to delayed diagnosis and appropriate treatment. At present, only three FDA-approved medications, adalimumab, secukinumab, and bimekizumab, are available for HS (*4, 5*). Nearly all other immunomodulatory medications are currently being used in an “off-label” manner, such as spironolactone and infliximab (*4, 5*). Many patients with advanced disease often fail multiple treatment modalities and are left with few options. Furthermore, the three FDA-approved therapies only show ∼50% improvement in disease in about 50% of study participants. Although these approved therapies reached clinical endpoints in their respective studies, their overall efficacies are still low compared to similarly approved drugs for other conditions such as psoriasis vulgaris. Therefore, there is a major unmet need for new therapies for HS, especially therapies that directly target the underlying pathophysiology of the disease and that can achieve better efficacy than existing agents.

The molecular complexity and multifactorial nature of HS pose a unique challenge for the development of effective therapies (*6*). Previous targeted treatments are focused on specific cytokines that are overexpressed in the blood and skin of most HS patients, but they do not consider individual genetic variability. This suggests the need for precision medicine approaches to identify novel therapeutic candidates and targets for HS beyond the current paradigm. Precision medicine is an emerging approach for disease treatment and prevention that takes into account individual variability in genes as well as other molecular measurements, leveraging large-scale datasets and big-data computational approaches. Development of novel therapeutics for HS has been hindered by several factors. There remains a large gap in knowledge regarding the pathophysiology and immunology of HS, including identifying targetable molecular pathways beyond those that are already known. Limited studies have recently shown that IL-1β, TNF-⍺, and IL12/23 are involved in the disease (*1*), as well as plasma cells (*7*), B cells (*7*), T-regulatory cells (*8*), and tertiary lymphoid structures (*9*). Anti-IL17 therapies including secukinumab and bimekizumab, have also recently been approved for HS (*10*). The advent of advanced technologies such as single-cell RNA sequencing (scRNAseq) and the public availability of -omics data in large cohorts in databases such as the Gene Expression Omnibus (GEO), provide a unique new opportunity to develop and apply computational integrative methods to refine the current knowledge about disease mechanisms, diagnostics, and therapeutics. Precision medicine approaches of multi-omic integration have worked successfully to identify therapeutic targets and combination therapy in other diseases (*11*) . Our group has pioneered and refined a systematic computational approach to predict novel therapeutics based on transcriptional signature reversal and have applied it to several diseases including breast cancer (*12*), Alzheimer’s disease (*11, 13*), inflammatory bowel disease (*14*), dermatomyositis (*15*), preterm birth (*16*), and COVID-19 (*17*). However, this methodology has not yet been systematically pursued for hidradenitis suppurativa or other inflammatory skin conditions.

Here, we propose an unbiased approach to the computational discovery of therapeutic candidates for hidradenitis suppurativa. Using a combination of existing locally generated and publicly available large-scale transcriptomic datasets of hidradenitis suppurativa and other inflammatory skin conditions, we identified immunologic signatures and pathways involved in the pathophysiology of disease (Figure 1A). We then leveraged an *in silico* computational drug repositioning approach to identify fulvestrant, pioglitazone, and sirolimus as novel therapies for HS based on reversal of gene expression signatures (Figure 1B). Finally, we validated the effects of fulvestrant, pioglitazone, and sirolimus using a novel preclinical *ex vivo* HS skin culture model (Figure 1C). No validated mouse model exists for HS, and although tissue allograft models have been studied, they poorly recapitulate the immunology of human skin. Therefore, we developed a model system for preclinical testing of computationally predicted therapies directly in human HS skin. We created an *ex vivo* model derived directly from HS skin from surgical excision of HS. Single cell suspensions of HS skin can be systematically perturbed with compounds to measure immune outputs such as inhibition of immune cell function and cytokine production, which can be extended to testing of single and combination drugs in a relatively high throughput manner. We expect this work to identify existing single and combination treatments that can be repurposed for the treatment of HS. Furthermore, we envision that this generalized systematic approach will be broadly applicable to the future discovery of drug candidates for other inflammatory skin diseases.

**Figure 1:**
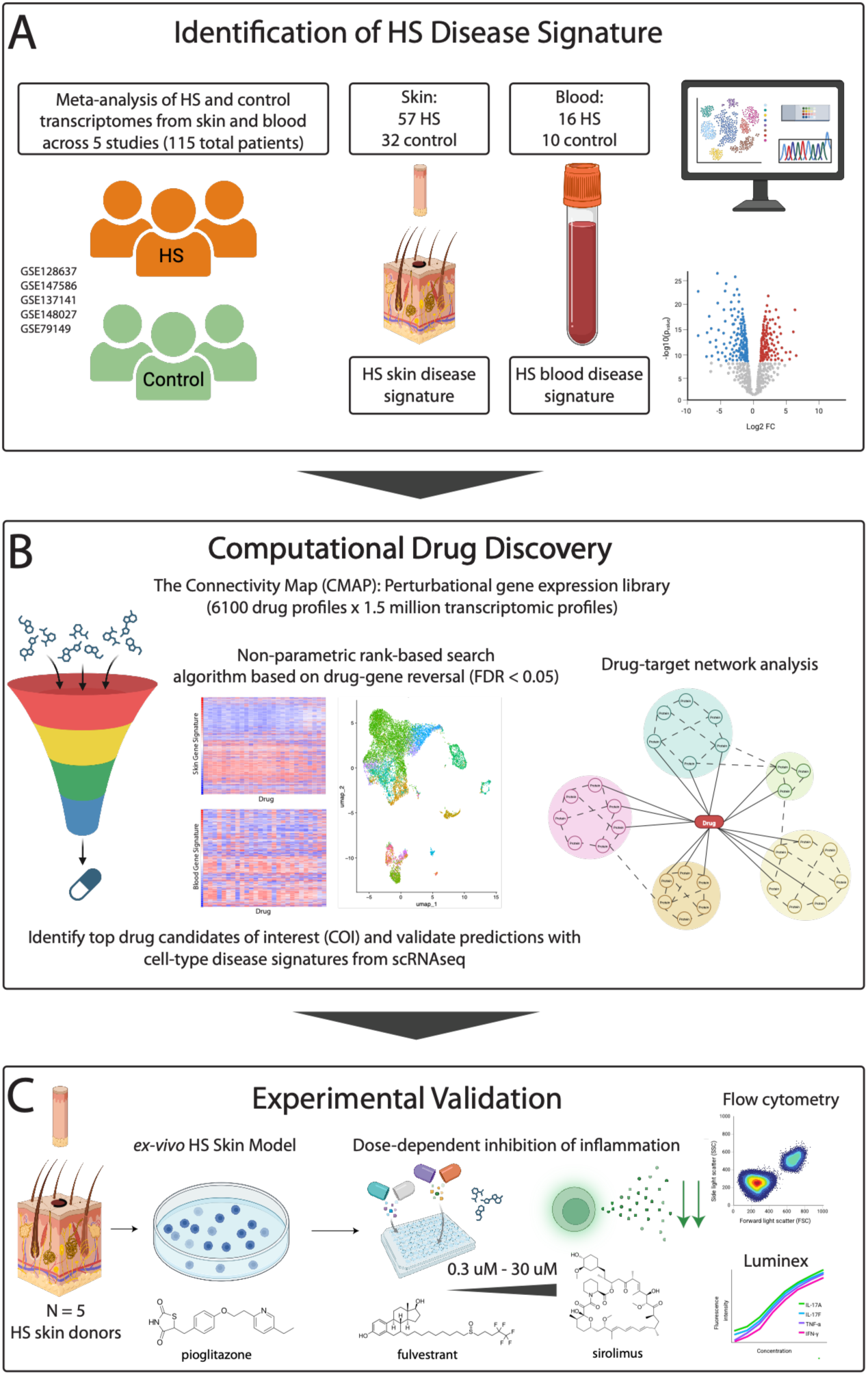
Computational discovery of novel therapeutics for hidradenitis suppurativa. Study overview. (A) Large-scale meta-analysis of transcriptomic datasets from HS patient skin and blood compared to healthy patients, and identification of an HS-specific disease signature. (B) Computational drug repurposing platform allowed for high throughput *in silico* screening of drug candidates based on reverse of disease-specific transcriptomic signatures and drug-target networks. (C) Experimental validation of predicted drug candidates using an *ex vivo* HS skin model via measurement of immune cell activation and cytokine production.

## Results

### Integrative meta-analysis of HS and healthy skin and blood identifies a targetable disease-signature

We conducted an integrative meta-analysis of microarray data from 57 samples of HS skin and 32 samples of skin from healthy controls across 4 studies, and 16 samples of HS blood and 10 samples of blood from healthy controls 1 study. In total there were 115 samples, 89 from HS patients and 42 from healthy controls (Figure 1A). Datasets were identified and downloaded from the GEO database using the keywords “hidradenitis suppurativa” and filtering for data taken from blood and from skin. The datasets used were GSE128637 (*18*) (10 HS, 11 control), GSE147586 (*19*) (6 HS, 5 control), GSE137141 (*20*) (16 HS, 8 control), GSE148027 (*21*) (25 HS, 8 control), and GSE79149 (*22*) (16 HS, 10 control). Using MetaIntegrator (*23*), we integrated the microarray datasets, allowing us to correct for batch effects. We identified a skin-specific transcriptomic signature of HS consisting of 3,165 upregulated genes and 3,499 downregulated genes (padj < 0.05, effect size >=0.5) (Figure 2A), and a blood-specific transcriptomic disease signature of HS consisting of 809 upregulated and 581 downregulated genes (padj < 0.05, effect size >=0.5, Figure 2B). Differentially expressed genes are shown in volcano plots, for HS skin vs. control skin (Figure 2A), and for HS blood vs. control blood (Figure 2B).

**Figure 2:**
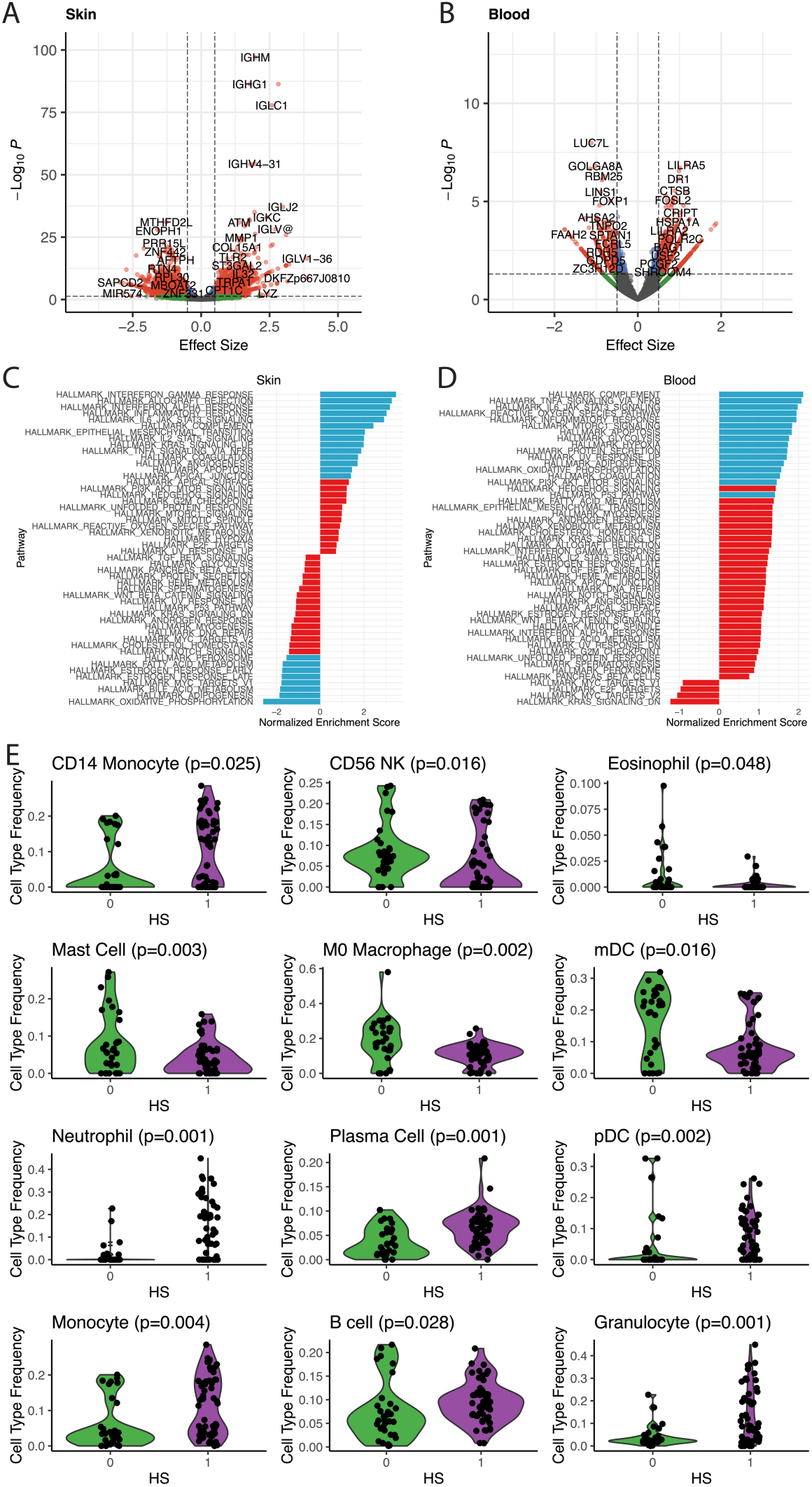
Integrative meta-analysis of HS and healthy skin and blood identifies a targetable disease-signature. Volcano plots for skin (A) and blood (B) disease signature. (C) and (D) Significant upregulation of IFN- γ, ⍺ (type I and type II interferon), and multiple JAK-STAT (IL6/JAK/STAT3, IL2/STAT5) pathways in skin and blood; down regulation of estrogen response, fatty acid metabolism, and oxidative phosphorylation. (E) Significant enrichment of monocytes, neutrophils/granulocytes, plasma cells, B cells, pDCs is observed in HS, while there is relatively downregulation of eosinophils, mast cells, and myeloid dendritic cells

The disease signature reliably differentiated HS from healthy controls in both tissues (Figure S1, Figure S2). Next, we conducted pathway analysis and cell-type deconvolution analysis to identify novel pathways underlying the pathophysiology of HS. We first conducted gene set enrichment analysis using FGSEA (*24*) with hallmark pathways to show that HS skin is enriched in upregulated pro-inflammatory pathways related to interferon-⍺, interferon- γ, JAK/STAT, IL-2, and NF-kB signaling. Downregulated pathways include those involved in estrogen response, adipogenesis, fatty acid metabolism, bile acid metabolism, and oxidative phosphorylation, suggesting dysregulation in hormonal and metabolic pathways in HS skin (Figure 2C). In HS blood, significant upregulated pathways include complement activation, TNF-⍺ signaling, IL6/JAK-STAT signaling, mTOR-signaling, and reactive oxygen species pathways (Figure 2D). Statistically significant normalized enrichments scores for each pathway are highlighted in blue, whereas non-significant pathways are shown in red. To identify cell types relevant in the pathogenesis of HS, we conducted cell-type deconvolution of the integrated dataset with ImmunoStates (*25*). We identified differential abundance of several immune cell types in HS skin compared to control skin. Statistically significant (FDR P-value < 0.05) increases were seen in monocytes, neutrophils, plasma cells, plasmacytoid dendritic cells, B-cells, and granulocytes, whereas decreases were seen in CD56 NK cells, eosinophils, mast cells, M0 macrophages, and myeloid dendritic cells in HS skin (Figure 2E).

### High throughput *in silico* screening identifies drug candidates predicted to reverse the global transcriptomic disease signature in HS

Using the disease and tissue-specific HS transcriptomic signatures from our integrated analysis, we then screened for drug candidates that were predicted to reverse the global disease signature of HS. Using the Connectivity Map (CMap) perturbational database, a drug expression database generated with human cancer cell lines, we create a computational pipeline to match gene expression profiles of disease vs. control with the gene expression profiles from FDA approved drugs (Figure 1B). The CMap contains 1.5 million gene expression profiles from ∼5,000 small-molecule compounds and ∼3,000 genetic reagents, representing a perturbational look-up table to make functional predictions. This algorithm comprises a non-parametric rank-based pattern-matching strategy to compare the differential gene expression signature for HS to differential gene expression drug (*12–16, 26*) . For the input gene signatures, there were 4001 genes in the total HS skin signature, and 1257 genes in the total HS blood signature. Drugs from the CMap were screened using this strategy, and a reversal score was calculated, with a negative score representing the predicted ability to reverse a disease gene signature towards healthy controls (Figure S3). Using a false discovery rate (FDR) of < 0.05, we identified a total of 109 drugs predicted to reverse the HS skin signature, and 260 drugs predicted to reverse the HS blood signature (Figure 3A, Figure S4, Figure S5). 27 predicted drugs were identified in both the blood and skin signature (Figure 3A). These drugs span diverse classes of medications, a subset of which are known immunomodulators. However, most of these drug candidates have unknown immunomodulatory potential and have not been systemically studied as potential treatments for HS. The major drug classes represented include traditional immunosuppressants (e.g. ciclosporin, sirolimus, methotrexate), non-steroidal anti-inflammatories (e.g. naproxen, meclofenamic acid), medications that target metabolic pathways (e.g. pioglitazone, rosiglitazone, troglitazone, metformin), antipsychotics (e.g. haloperidol, fluphenazine, clozapine, thioridazine), antiepileptics (e.g. valproic acid), and hormonal therapies (e.g. estradiol, letrozole, fulvestrant). We calculated the gene expression profiles and reversal scores of each of the 27 hits present in both blood and skin, and all of which were predicted to strongly reverse the disease signature (Figure 3B, 3C). Red denotes upregulation of a specific gene whereas blue denotes downregulation of a specific gene. The strength of reversal (range -1 to 1 with more negative equating to a stronger predicted reversal) was plotted on a heatmap for each of the 27 drugs, which allowed us to select candidate drugs that were strongly predicted to reverse the HS disease signature in both tissues (Figure 3D).

**Figure 3:**
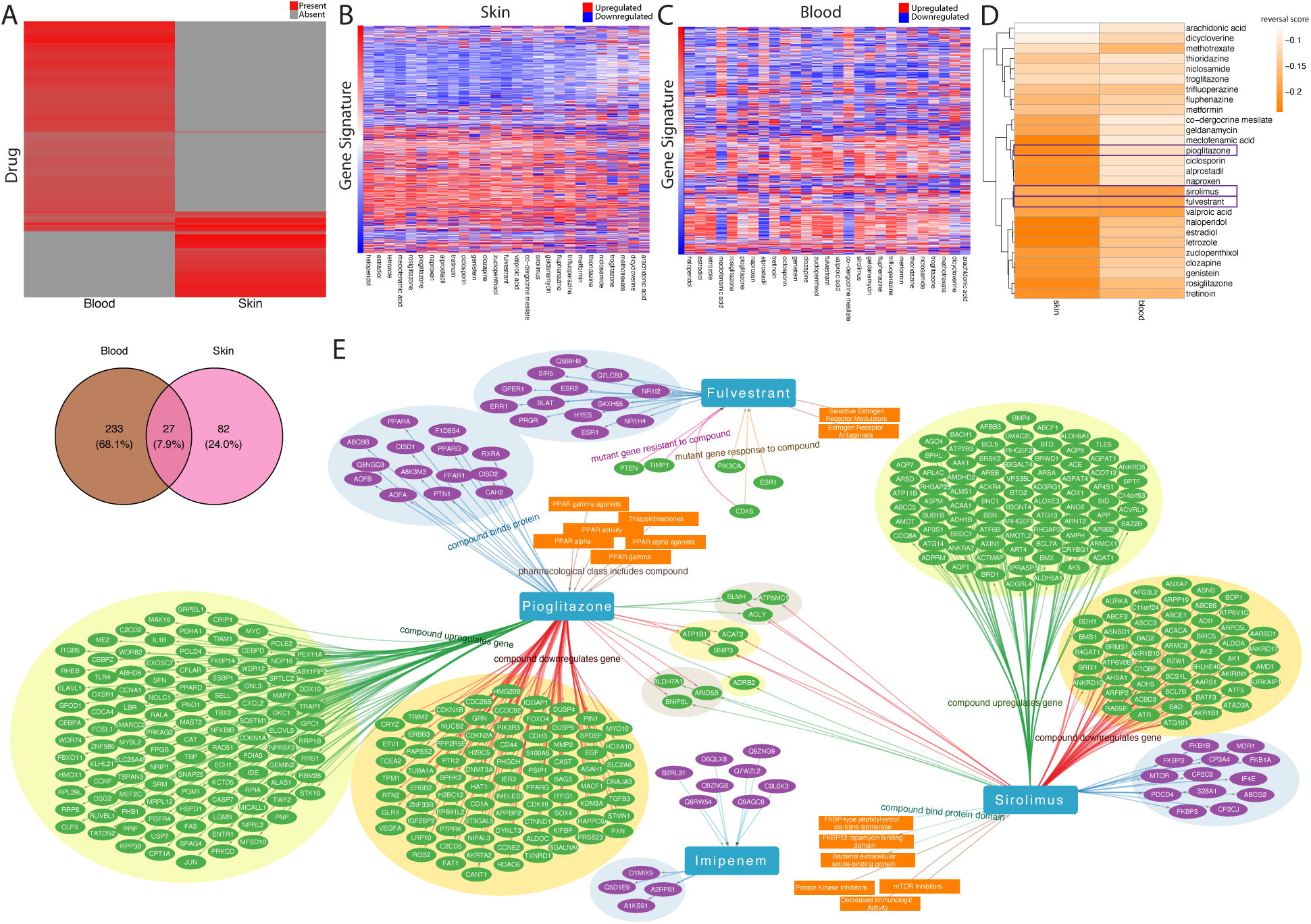
High throughput *in silico* screening identifies drug candidates predicted to reverse the global transcriptomic disease signature in HS. (A) 109 drugs identified from skin 260 drugs identified from blood, with 27 overlapping. (B) and (C) gene drug reversal signatures for the 27 drugs found in both skin and blood. There are 4001 genes in total gene signature for skin, and 1257 genes in total gene signature for blood. (D) reversal scores for skin and blood for 27 drugs. (E) Drug-target network from 4 selected drugs

We selected three drugs for further comparison using drug-target networks (fulvestrant, pioglitazone, sirolimus) due to their predicted ability to reverse both the HS skin and blood disease signatures. Sirolimus is a known immunosuppressant (*27*) targeting the mammalian target of rapamycin (mTOR) that is primarily used for the prevention of transplant rejection, although has not been systematically tested in HS. Pioglitazone is a thiazolidinedione class peroxisome proliferator-activated receptor gamma (PPAR-γ) agonist (*28*) that is FDA approved for diabetes mellitus but has not been studied as a treatment for HS. Finally, fulvestrant is a selective estrogen receptor modulator and antagonist (*29*) that is FDA approved for hormone receptor-positive breast cancer but has not been studied as a treatment for inflammatory skin disease or HS. We also selected imipenem for comparison, a carbapenem class antibiotic, as it represents a structural analogue of ertapenem which is commonly used as an intravenous rescue therapy for HS flares but with an unknown mechanism of action in humans (*30*) .

Imipenem was identified as a top hit in the blood signature of HS but was not one of the top hits in the skin. A drug-target network using the SPOKE knowledge network (*31*) and DrugBank (*32*) was generated to identify relationships between drugs, mechanisms, and targets (Figure 3E, Figure S6). We found that fulvestrant was able to modulate 13 human protein targets, beyond the canonical estrogen receptor. Pioglitazone and sirolimus had a significantly larger number of protein targets. Interestingly, fulvestrant and pioglitazone, and fulvestrant and sirolimus do not have specific overlapping protein targets, but pioglitazone and sirolimus have 10 shared targets. Imipenem as expected only had known binding to bacterial proteins, consistent with its known mechanism as a beta-lactam antibiotic (Figure 3E). This shows that the identified drug candidates are predicted to modulate the disease signature of HS through previously unknown pathways.

### Drug candidates target the HS disease signature in skin immune cells relevant to HS pathogenesis

Next, we used single cell RNA sequencing data to validate the predicted drugs from our meta-analysis. We hypothesized that the predicted drugs from our integrated microarray dataset could potentially target specific skin immune cell types involved in the pathogenesis of HS. To identify and validate our hypothesis of whether predicted COIs could reverse the disease signature in a specific cell type, we queried a dataset of single cell RNA sequencing conducted on HS skin and on healthy skin (GSE155850) (*8*). We visualized scRNAseq data obtained from >20,000 sorted immune cells taken from two samples of HS skin and integrated with data from two samples of normal skin (Figure 4A, 4B). Data integration was conducted with Seurat (*33*). Clusters were identified based on unsupervised clustering with UMAP, and cell type identification was conducted using label transfer from previously annotated scRNAseq datasets of human skin immune cells (*34, 35*). Cluster comparison was conducted for the integrated scRNAseq data. Granulocytes were not included in this analysis. We identified 13 cell types from cell type annotation that are present in significant abundance across control and HS: Dendritic cells (DC1, DC2, DC3, moDC2), macrophages (Mac1, Mac2), monocytes (Mono), inflammatory monocytes (InfMono), B cells (B), natural killer cells (NK), innate lymphoid cells (ILC_NK), T-resident memory cells (Trm3), and T-migratory memory cells (Tmm3) (Figure S7). The Trm3 and Tmm3 groups contain both CD4 and CD8+ T cells (Figure 4A, 4B). Next, we conducted differential gene expression from each of the skin immune cell types using Deseq2, comparing HS to control skin, to identify cell-type specific transcriptomic signatures in HS (Figure 4C). Gene set enrichment analysis was conducted for each cell type to identify specific enriched molecular pathways (Figure 4D). Several cell types exhibited significant upregulation of IFN-alpha and IFN-gamma pathways, including DC1, Mac2, InfMono, ILC-NK, and Mono. There was significant downregulation of estrogen response and androgen response pathways in DC1, Mac2, and InfMono, suggesting a role for modulation of hormonal pathways in these cell types. We then inputted each cell type-specific HS disease transcriptomic signature into the computational drug discovery pipeline we previously used to identify therapies from the integrated microarray datasets. We calculated top predicted drugs for each cell type. With an FDR < 0.05, we identified 135 hits for B cells, 93 hits for DC1, 83 hits for DC2, 48 hits for DC3, 61 hits for ILC_NK, 71 hits for InfMono, 142 hits for Mac1, 236 hits for Mac2, 71 hits for moDC2, 159 hits for Mono, 79 hits for NK, 138 hits for Tmm3, and 98 hits for Trm3. We asked whether the four drugs identified previously from the global HS disease signature, sirolimus, fulvestrant, pioglitazone, and imipenem, could be predicted to reverse the HS disease signature in specific cell types. Interestingly, several of the COIs identified from our microarray analysis were predicted to also reverse the HS disease signature. Of the 27 drugs identified from the integrative microarray dataset, all 27 appeared as a hit in at least 1 or more cell types. Sirolimus and fulvestrant were hits in all 13 cell types. Pioglitazone and imipenem were also predicted to reverse the HS disease signature, though in a smaller subset of cell types. Pioglitazone was a hit in 5 cell types and imipenem was a hit in 2 cell types (Figure 4E).

**Figure 4:**
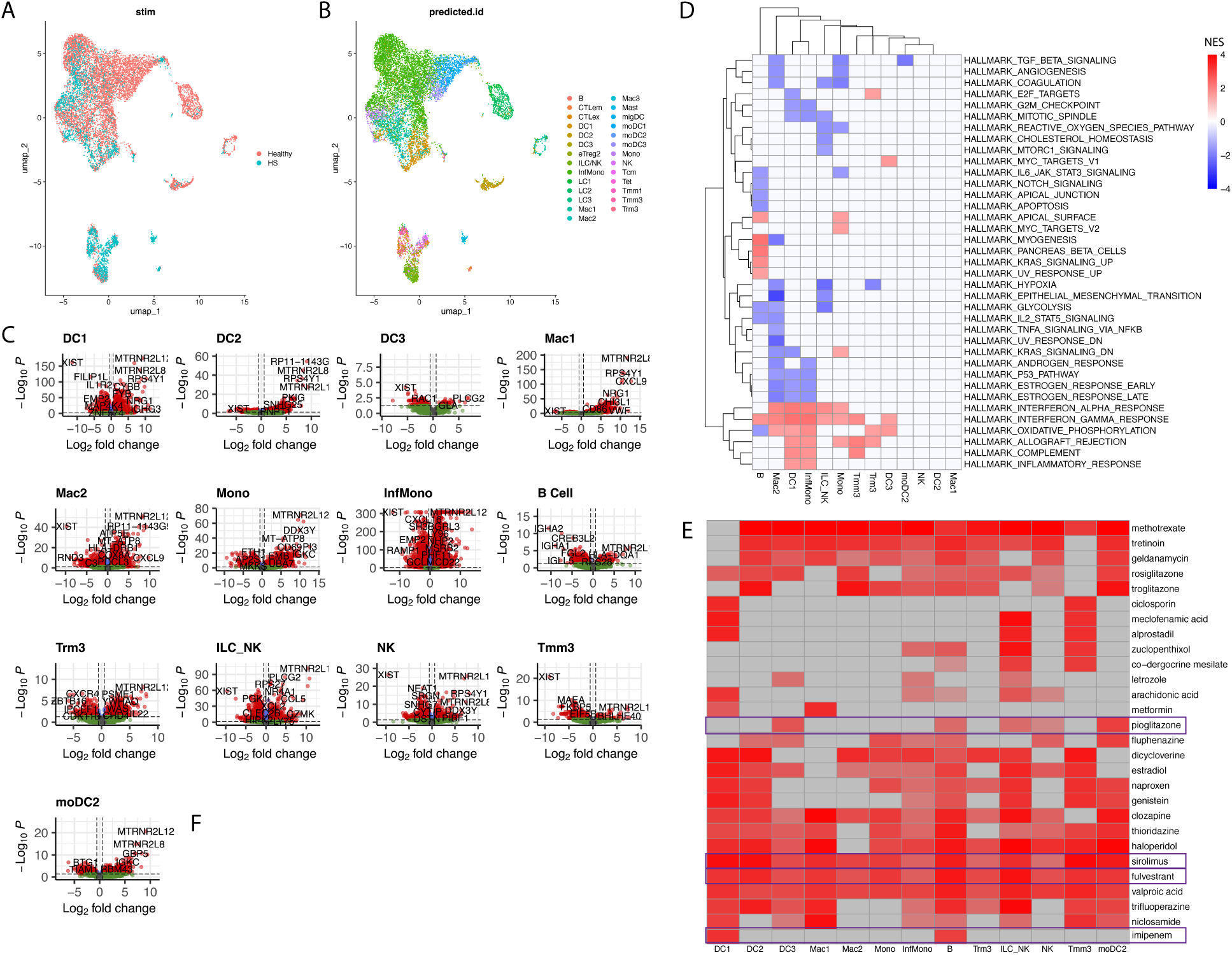
Drug candidates target the HS disease signature in skin immune cells relevant to HS pathogenesis. (A) UMAP of combined scRNAseq data from skin immune cells HS vs. healthy control, as well as cell type annotation of each skin immune cell type (B). (C) Volcano plots of each cell type differential gene expression analysis (padj < 0.05, log2FC > 0.5). (D) Pathway analysis identified cell-type specific pathway upregulation and downregulation in HS skin. IFN response for both ⍺ and γ is particularly upregulated in dendritic cells, macrophages, monocytes, ILC, and B cells and T cells. (E) Computational drug screening showing hits across 13 cell types. Sirolimus and fulvestrant were present as hits in all 13 cell types. Pioglitazone and imipenem were present in the predictions in a smaller subset of cell types.

### Predicted drug candidates inhibit T-cell proliferation and activation and block pro-inflammatory cytokine production in an *ex vivo* HS skin model

We selected four compounds for testing in the *ex vivo* HS skin model: fulvestrant, pioglitazone, sirolimus, and imipenem. These compounds were chosen for several reasons. Both fulvestrant and sirolimus were predicted to reverse the disease signature of HS in all 13 cell types identified, as well as in the global HS disease signature from our meta-analysis.

Sirolimus is a known immunomodulator and transplant medication, but has not been studied systematically in HS beyond a single case series (*36*) . Both fulvestrant (*29*) and pioglitazone (*28*) represent novel mechanisms of action of HS treatment, without prior significant direct validated evidence of immunomodulatory activity in HS skin. We hypothesized that modulation of hormonal pathways and immunomodulatory pathways could alleviate inflammation in HS skin. Finally, imipenem was chosen as its carbapenem analogue ertapenem has previously been used in humans as a rescue therapy for severe HS, but its immunomodulatory activity on host function is unclear (*30*).

To experimentally validate these predicted drug candidates, we needed to develop a framework for preclinical testing. No validated mouse model exists for HS, and although tissue allograft models have been studied, they poorly recapitulate the immunology of human skin. Therefore, we developed a model system for preclinical testing of computationally predicted therapies using an *ex vivo* HS skin model assay. We utilized excisional specimens from HS patients undergoing surgery of active HS lesions and digested the tissue into single-cell suspensions enzymatically, based on previous work from our group (*8, 37*) . Specific immune cell types were assessed using flow cytometry, including CD4 T cells, CD8 T cells, and T-regulatory cells (Tregs). We collected skin samples from five different human donors and experiments were carried out in triplicate. Samples were aliquoted into U-bottom plates and cultured for 2-3 days. Compounds were incubated with HS skin derived immune cells in a dose dependent manner ranging from 0 uM (vehicle) to 30 uM, which represents a physiological range of concentrations for these drugs based on previous studies. Flow cytometry was used to measure immune cell viability, proliferation, and activation. We found that all four drugs did not have a significant effect on cell viability (Figure 5A), but that fulvestrant, pioglitazone, and sirolimus significantly inhibited proliferation of CD4 T cells, CD8 T cells, and Tregs in a dose dependent fashion (Figure 5B-D) (*p<0.05, **p<0.01). Next, we used multiplexed Luminex cytokine assays to measure the proteomic impact of these predicted drugs on the production of proinflammatory cytokines relevant to HS pathogenesis. Supernatants were taken from HS cells following incubation with the predicted drugs. Cytokines of interest quantified from the supernatant include IL-17A, IL17-F, G-CSF, IFN- γ, CXCL10, TNF-⍺, CCL3, GM-CSF, IL-1beta, IL-6, and IL-8. Normalized fluorescence intensities from Luminex immunoprofiling converted into cytokine concentrations using a standard curve. We found that sirolimus, pioglitazone, and fulvestrant (Figure 5E), but not imipenem, suppressed proinflammatory cytokine production from HS skin, including IL-17A/F, IFN-γ, CXCL10, and G-CSF in a dose dependent manner compared to vehicle.

**Figure 5:**
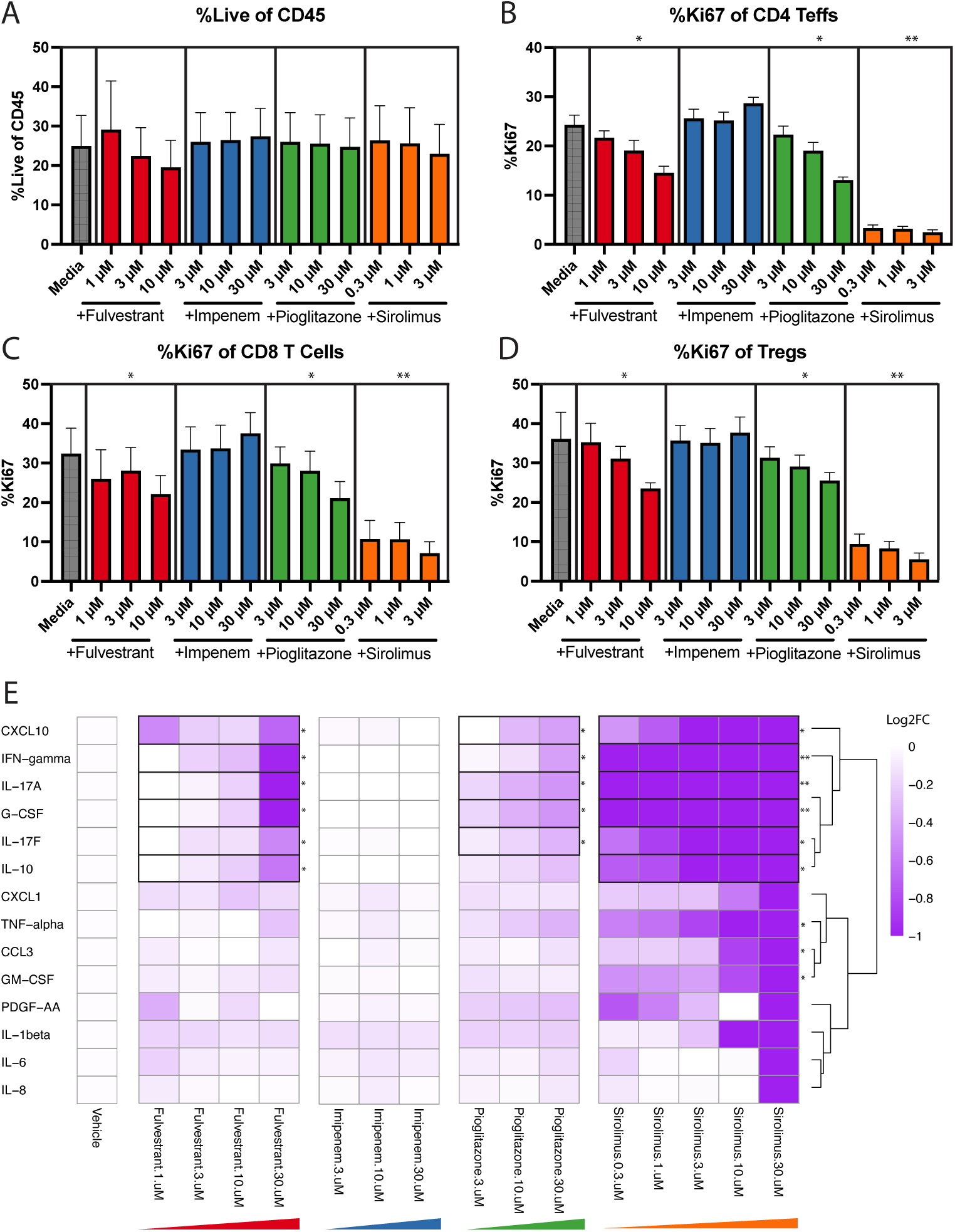
Predicted drug candidates inhibit T-cell proliferation and activation and inhibit pro-inflammatory cytokine production in an *ex vivo* HS skin model. Flow cytometry analysis from skin-derived immune cells relevant to HS pathogenesis. (A) % of cells that remain viable. Proliferation rate of specific T cell subtypes including CD4 (B), CD8 (C), and Tregs (D). Significant dose dependent downregulation of proliferation of CD4, CD8, and Tregs by fulvestrant, pioglitazone, and sirolimus are observed (*p < 0.05, **p < 0.01). (E) Normalized fluorescence intensities from Luminex immunoprofiling converted into cytokine concentrations using a standard curve, reported as log 2-fold change (*p < 0.05, **p < 0.01). Fulvestrant, pioglitazone, and sirolimus suppressed proinflammatory cytokine production from HS skin, including IL-17A/F, IFN-γ, CXCL10, and G-CSF in a dose dependent manner compared to vehicle.

## Discussion

Hidradenitis suppurativa is a severe chronic inflammatory disease for which precision medicine is urgently needed. Only three advanced FDA approved treatments for HS exist, which are monoclonal antibodies that target TNF-⍺ (adalimumab) and IL-17 (bimekizumab and secukinumab). Although these therapies demonstrate some efficacy, patients often relapse and exhibit refractory disease. Furthermore, they are only approved for moderate and severe disease, at which point fibrosis and tunnel formation has already taken place. Interestingly, only about ∼50% of trial participants achieve the FDA-accepted clinical endpoint of achieving a 50% reduction of inflammatory lesions. Higher efficacy is needed, as many patients still have a significant remaining disease burden despite treatment with these approved therapies.

Therefore, there is a major unmet need for new therapies for HS that directly target the underlying pathophysiology of the disease and that can achieve better efficacy than existing agents. Given the need for safe and more efficacious disease-modifying treatments, computational drug repurposing has gained interest due to its faster development, lower costs, and improved safety since these drugs have already been FDA approved for other indications. Advancements in the generation of large-scale perturbational drug databases offer new opportunities for discovering promising drug candidates for diseases with few treatment options. High-throughput *in silico* drug screening approaches have identified numerous repurposed candidates for other inflammatory diseases in the past.

In this study, we proposed the repurposing of three existing FDA approved drugs: fulvestrant (estrogen receptor modulator approved for breast cancer), pioglitazone (PPAR-γ agonist approved for type 2 diabetes mellitus), and sirolimus (mTOR inhibitor used as prophylaxis for transplant rejection), to reverse disease and cell-type specific gene expression alterations across skin and blood in HS. Our drug-screening strategy is driven entirely by human data taken from skin and blood, comprising large-scale genomic datasets, drug perturbation libraries generated in human cell lines, and preclinical validation in an *ex vivo* model of HS skin to maximize clinical translation. We demonstrated that treatment of HS skin with fulvestrant, pioglitazone, and sirolimus inhibited immune cell activation and proliferation and reduced the production of HS-specific proinflammatory cytokines in a dose-dependent manner, positioning these drug candidates for rapid testing in humans in randomized clinical trials. This successful approach combining computational drug discovery based on large-scale transcriptomics data and preclinical validation in a translational model of HS skin highlights the power of precision medicine approaches in the discovery of new therapies to target dysregulated molecular signatures in HS. Our integrative computational approach demonstrated here will be broadly generalizable to other chronic inflammatory diseases beyond HS that require precision medicine approaches, such as systemic lupus erythematosus, scleroderma, and inflammatory bowel disease.

The identification of fulvestrant, pioglitazone, and sirolimus as novel therapies for HS has far-reaching implications. Early treatment of HS has typically focused on topical therapy with clindamycin and benzoyl peroxide and systemic antibiotics such as doxycycline, clindamycin, and rifampin. Other adjunctive therapies include spironolactone and intralesional corticosteroid injections. Off-label treatments for severe disease include infliximab (anti-TNF) infusions and intravenous ertapenem (*4, 5*) . What’s clear is that no existing treatments for HS target dysregulation in immunometabolic pathways. Significant research has shown that HS is comorbid with metabolic syndrome, which is also associated with chronic inflammation. Our findings suggest that modulation PPAR-γ with drugs such as pioglitazone may represent a novel viable approach in treating HS. Previously, the immunologic effects of pioglitazone have been studied in multiple sclerosis (*38*) . In an orthogonal direction, modulation of the estrogen receptor has received interest as a potential immune target. A few reports have demonstrated a potential immunomodulatory role for fulvestrant, but direct evidence is sparse (*39, 40*) . The estrogen receptor is expressed in numerous cell types, including immune cells (*41–43*), suggesting a potential hidden role of immuno-hormonal pathways in the pathogenesis of HS. Sirolimus is a known immunosuppressant, so at first glance it may not be surprising that sirolimus is able to inhibit inflammation in HS. Interestingly, compounds with similar mechanisms of action such as tacrolimus or everolimus were not identified, suggesting that sirolimus may better target the dysregulated gene networks in HS. Sirolimus is a potential candidate as well for topical therapy for HS, as topical formulations are now FDA approved for other indications (facial angiofibromas) (*44*) . Further clinical trials will need to be conducted to test the efficacy of our proposed therapies in humans.

Several limitations of this study should be acknowledged. First, although our computational framework leverages large-scale human transcriptomic datasets derived from skin and blood, some of this data is based on microarrays, which may obscure important cell-type–specific effects and rare immune populations relevant to HS pathogenesis. While we partially mitigate this through cell-type–resolved analyses and functional validation in an ex vivo HS skin model, future studies incorporating single-cell and spatial transcriptomics will be necessary to further refine drug–cell interactions and therapeutic targeting, especially drug targeting to spatially-resolved locations in the skin. Second, drug perturbation signatures were derived from established human cell lines, which may not fully recapitulate the complex cellular microenvironment, hormonal influences, and immune–stromal interactions present in HS skin *in vivo*. Third, although *ex vivo* skin cultures provide a translationally relevant human tissue platform, they cannot model long-term disease evolution, fibrosis, tunneling, or systemic immune contributions that characterize chronic HS. Additionally, dosing and exposure in ex vivo systems may not directly translate to pharmacokinetics, safety, or efficacy in patients. We address this in our preclinical studies by using clinically relevant concentrations that are realistically found *in vivo*. Finally, while our study identifies mechanistically unique FDA-approved drug candidates that have known clinical safety and tolerability, clinical efficacy must still be tested systematically in prospective, randomized controlled clinical trials of HS patients, which is outside the scope of this study but the next logical step for future work. Despite these limitations, our integrative human-data–driven -omics approach provides a strong foundation for precision therapeutic discovery in HS and other chronic inflammatory diseases.

In summary, this study introduces a novel approach to drug discovery for hidradenitis suppurativa by combining integrative genomics on data derived directly from diseased tissue, computational drug repositioning using large perturbational databases, and validation in an *ex vivo* HS skin model. Our methodology provides a robust framework for identifying effective therapies through real-world evidence. Our preclinical studies demonstrate the ability of our drug candidates to directly inhibit cell types involved in the pathogenesis of HS and abrogate production of proinflammatory cytokines. Our findings demonstrate the ability the predicted drugs to reverse transcriptomic networks that are dysregulated in HS, paving the way for precision medicine by leveraging artificial intelligence and large -omics datasets (*45, 46*) .

## Materials and Methods

### Transcriptomic meta-analysis

For the meta-analysis of gene expression datasets, we employed the MetaIntegrator framework (*23, 25*) , an established computational method specifically designed to integrate and analyze transcriptomic data from multiple independent studies. Meta-analysis of gene expression data is crucial in consolidating findings across different datasets, enabling the identification of robust and reproducible gene signatures associated with the condition of interest, even in the presence of biological and technical variability. MetaIntegrator calculates effect sizes for each gene across all datasets, using the standardized mean differences (SMDs) to quantify the magnitude and direction of association between gene expression and disease state:

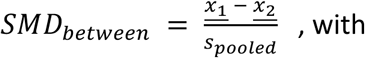

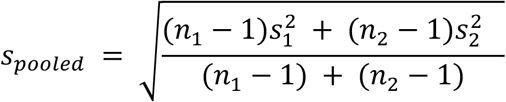

where <u>𝑥</u>_1_ is the mean of group 1, <u>𝑥</u>_2_ is the mean of group 2, 𝑛_1_ is the number of observations in group 1, 𝑛_2_ is the number of observations in group 2, 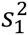 is the variance of group 1 and 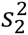 is the variance of group 2. *s_pooled_* is therefore the pooled standard deviation, which is a weighted average of the standard deviations of the two groups. To address inter-study heterogeneity—a common issue in meta-analysis—the method uses a random-effects model. This statistical approach assumes that effect sizes may vary across studies, accounting for both within- and between-study variability. The random-effects model thus provides a more generalizable estimate of effect size, reducing the risk of bias that might arise from directly pooling data. Gene signatures are then identified based on combined effect sizes and significance thresholds, prioritizing genes with consistent associations across studies. This rigorous approach provides a robust and reproducible set of candidate genes for further validation and exploration, particularly valuable for complex diseases where single-cohort studies may lack sufficient power.

### Pathway analyses

For pathway analyses to identify immune pathways and signatures associated with HS, we inpuued the sorted differentially expressed gene lists defining the log 2 fold change and the adjusted p-values into the Bioconductor package fgsea (v 1.10.1) for gene set enrichment analysis (GSEA) (*20*) with Hallmark pathways taken from the MSigDB (*47*).

### Cell-mixture deconvolution analyses

The ImmunoStates (*25*) package was used to carry out cell type deconvolution analysis. Cell-mixture deconvolution is a computational approach designed to infer the proportions of distinct cell types present within a bulk tissue sample. This method enables the dissection of complex biological samples into their cellular components, offering a deeper understanding of the sample’s cellular heterogeneity. For this analysis, we utilized the ImmunoStates basis matrix, a reference framework comprising gene expression profiles for well-characterized immune cell types, including T cells, B cells, monocytes, and natural killer cells, among others. The deconvolution process leverages a set of marker genes that are selectively expressed by specific cell types. By comparing the expression levels of these markers in the sample to those in the reference basis matrix, a linear modeling framework is applied to estimate the relative proportions of each cell type. This quantitative inference is crucial for studying immune-related diseases and for understanding how shifts in cellular composition may reflect pathological states or therapeutic responses. Importantly, this method enables the analysis of bulk RNA-sequencing data, which represents the averaged expression across all cells in a sample, and resolves it into meaningful contributions from individual cell types. By using the ImmunoStates matrix, we tailored our analysis to focus on immune cells, offering insights into the immune landscape of the disease. This approach not only enhances our ability to interpret gene expression data but also provides a critical foundation for identifying potential biomarkers, exploring disease mechanisms, and developing targeted therapies. The false discovery rate was controlled by the Benjamini–Hochberg method.

### Single Cell RNA Sequencing Analysis

Single cell RNA sequencing data was obtained from our previous work (GSE155850) (*8*). Briefly, skin immune cells (Live, singlet, CD3-, CD19- events) were sort-purified, counted, and loaded onto a 10x Single Cell 3’v3 GEMS chip and sequenced by the 10x Genomics Core (UCSF). Fastq files were aligned to GRCh38 with Cell Ranger version 3.0.2. Clustering and differential expression analysis was performed using Seurat (*33*) . Data was integrated and scaled, UMAP/PCA was run, and clusters were identified. Cell type annotation was completed using label transfer from previously annotated scRNAseq datasets of human skin immune cells (*34, 35*) . We identified cell types from cell type annotation that are present in significant abundance across control and HS skin. Differential gene expression for each cluster was conducted using Deseq2. Differential expression cell-type specific disease signatures were then combined with the computational drug repurposing pipeline to identify predicted drugs, as described below.

### Computational drug repurposing pipeline

Inspired by the Kolmogorov-Smirnov statistic and the framework introduced by Lamb et al. (*48, 49*) in the original CMap database, the pipeline employs a nonparametric rank-based method to evaluate the concordance between disease-associated gene expression profiles and drug-induced expression signatures. A prefiltering step, adapted from prior drug repurposing methodologies, was applied to refine the input data. Genome-wide differential expression profiles from the CMap database were then analyzed, and candidate drugs were ranked based on the statistical significance of their reverse correlation with the preeclampsia signature, with adjustments for multiple testing. For drugs tested under varying conditions, the profile with the strongest reversal was selected for each compound.

### Network Analysis

To better understand the potential relationship between the top drug and preeclampsia, we explored relationships between genes, compounds, and related diseases via the SPOKE knowledge network (*31*) which includes information from DrugBank (*32*), ChEMBL, and Gene Ontology. To characterize the top drug candidates from the top HS signature from both skin and blood, we utilized the SPOKE knowledge network, which includes relationship information from DrugCentral, Chembl, BindingDB, and CMAP/LINCS databases, to identify nearest neighbors and visualize drug relationships in the network. The nearest neighbor networks were determined using nodes of class “Compound” (including the drugs of interest), “Gene”, “Pharmacological Class”, and “Protein” or “Protein Domain”. Edges included “Protein Transports Compound”, “Compound Binds Protein”, “Compound is a Compound”, “Compound upregulates/downregulates Gene”, “Compound has role as Compound”, “Pharmacological Class includes Compound”, and “mutant Gene respond/resistant to Compound”. In order to understand the drug-disease relationships, we utilized the shortest paths in the SPOKE knowledge networks between hidradenitis suppurativa (DOID:2282) or hidradenitis (DOID: 2280) and each HS drug candidate. Additional nodes include “Disease” and edge include “Compound treats Disease” where edges are filtered to those being studied in at least Phase 3 clinical trials or beyond.

### Ethics Statement

Discarded de-identified tissue was certified as Not Human Subjects Research per institutional guidelines. Patients providing tissue with associated clinical metadata provided written, informed consent under protocol 13-11307. The UCSF Institutional Review Board approved the proposed studies (approval 16-19770).

### Compounds

All compounds were resuspended within the ex-vivo assay medium. Sirolimus, pioglitazone, fulvestrant, and imipenem (Sigma-Aldrich) were prepared at a 1000× concentration in the solvents listed below and diluted to targeted concentrations based on experiments with the *ex-vivo* HS skin model.

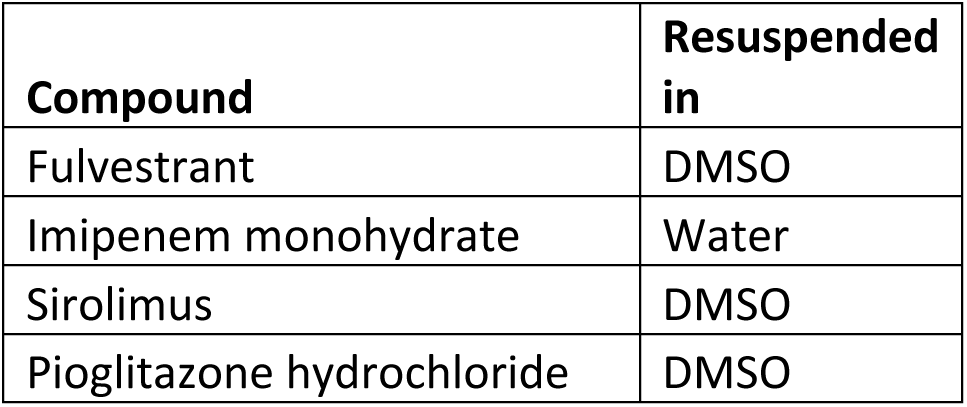

| Compound | Resuspended in |
| --- | --- |
| Fulvestrant | DMSO |
| Imipenem monohydrate | Water |
| Sirolimus | DMSO |
| Pioglitazone hydrochloride | DMSO |

### Ex-vivo HS skin model

To produce single-cell suspensions from HS skin (*37*), HS explants were enzymatically digested directly following incubation using collagenase IV (1000 U/mL, Worthington, LS004186) and DNAse (200 μg/mL) for 2 h in RPMI +10% FBS. Specimens were cryopreserved post-processing and thawed prior to the assay. Samples were then plated in a 96-well U-bottom plate, with 300 000 cells allocated to each well, and cultured for 2–3 days at 37°C in a 5% CO 2 incubator.

### Flow cytometry

Following *ex vivo* culture, samples were then stained extracellularly using Ghost Dye Violet 510 (Tonbo Bioscience, 13- 0870-T100), Brilliant Violet 711 anti-human CD45 Antibody (BioLegend, 304050), PE- CF594 Mouse anti-human CD4 (BD Horizon, 562281), CD3 Monoclonal Antibody APC-eFluor 780 (Invitrogen, 47- 0036- 42), Brilliant Violet 650 anti-human CD8a Antibody (BioLegend, 301042), CD127 PE Clone hIL7R-M21 Cat 557938, CD19 Alexa700 CloneHIB19;Cat:302225, ICOS BV785 Clone 313533 Cat: C398.4A and BD APC Mouse Anti-Human CD25 (BD Biosciences, 340939). Fixation and permeabilization was performed using the eBioscience Intracellular Fixation & Permeabilization Buffer Set (Invitrogen, 88-8824-00). Intracellular staining was performed using the FOXP3 Monoclonal Antibody, eFluor 450 (Invitrogen, 48- 4776- 42), anti-IL1R2 FITC Clone 34141; Cat: MA5-23662, anti-CTLA4 PerCPe710 Clone 14D3 Cat: 46-1529-42, and anti-Ki67 PECy7 Clone B56 Cat 561283. Samples were then run on an LSRFortessa X-20 Cell Analyser (BD Biosciences, Milpitas, CA) and analysed using FlowJo v10.10 Software (BD Life Sciences, Franklin Lakes, NJ).

### Cytokine release assays

Culture supernatants were collected and frozen at -80. Cytokine concentrations were then quantified via the Human Cytokine Panel A 48-Plex Discovery Assay Tissue Homogenate (Luminex, Eve Technologies, Calgary, AB, Canada).

### Statistics and reproducibility

The statistical analyses for each part of the approach are described above. A threshold of P = 0.05 for false discovery rate and for adjusted P-values was set. The code is publicly available to ensure the reproducibility of our results.

## List of Supplementary Materials

Figure S1: PCA of datasets used in meta-analysis

Figure S2: Meta-analysis scores differentiating HS from control skin

Figure S3: Histogram distribution of drug reversal scores

Figure S4: Aggregate gene-drug reversal signatures for HS skin

Figure S5: Aggregate gene-drug reversal signatures for HS blood

Figure S5: Drug-protein target interactome

Figure S6: Cell type abundances for scRNAseq annotation

Table S1: Drug-disease reversal scores for the HS skin signature

Table S2: Drug-disease reversal scores for the HS blood signature

Table S3: Reversal scores for drugs identified in both skin and blood signatures

## Acknowledgments

We gratefully acknowledge the hidradenitis suppurativa patients who generously contributed tissue samples to support this research.

## Author contributions

All authors were involved in drafting the article or revising it critically for important intellectual content, and all authors approved the final version to be published.

## Data and materials availability

The code and data that support the findings of this study are available from the corresponding author upon reasonable request.

