## Supplemental Information for "Computational discovery of precision therapeutics for hidradenitis suppurativa"

**Affiliations:**
[1] Department of Dermatology, University of California, San Francisco, San Francisco, CA, United States
[2] Department of Pediatrics, University of California, San Francisco, San Francisco, CA, United States
[3] Bakar Computational Health Sciences Institute, University of California, San Francisco, San Francisco, CA, United States
[4] immunoLogic, San Francisco, CA, United States
[5] Department of Dermatology, University of Connecticut, Farmington, CT, United States
[6] Department of Surgery, UCSF, San Francisco, California, United States

***Corresponding Authors:**
Ernest Y. Lee
Department of Dermatology, University of California, San Francisco, San Francisco, CA, United States
Bakar Computational Health Sciences Institute, University of California, San Francisco, San Francisco, CA, United States

Margaret Lowe
Department of Dermatology, University of California, San Francisco, San Francisco, CA, United States

Marina Sirota
Bakar Computational Health Sciences Institute, University of California, San Francisco, San Francisco, CA, United States

### Supplemental Figures:

### Figure S1: PCA of datasets used in meta-analysis

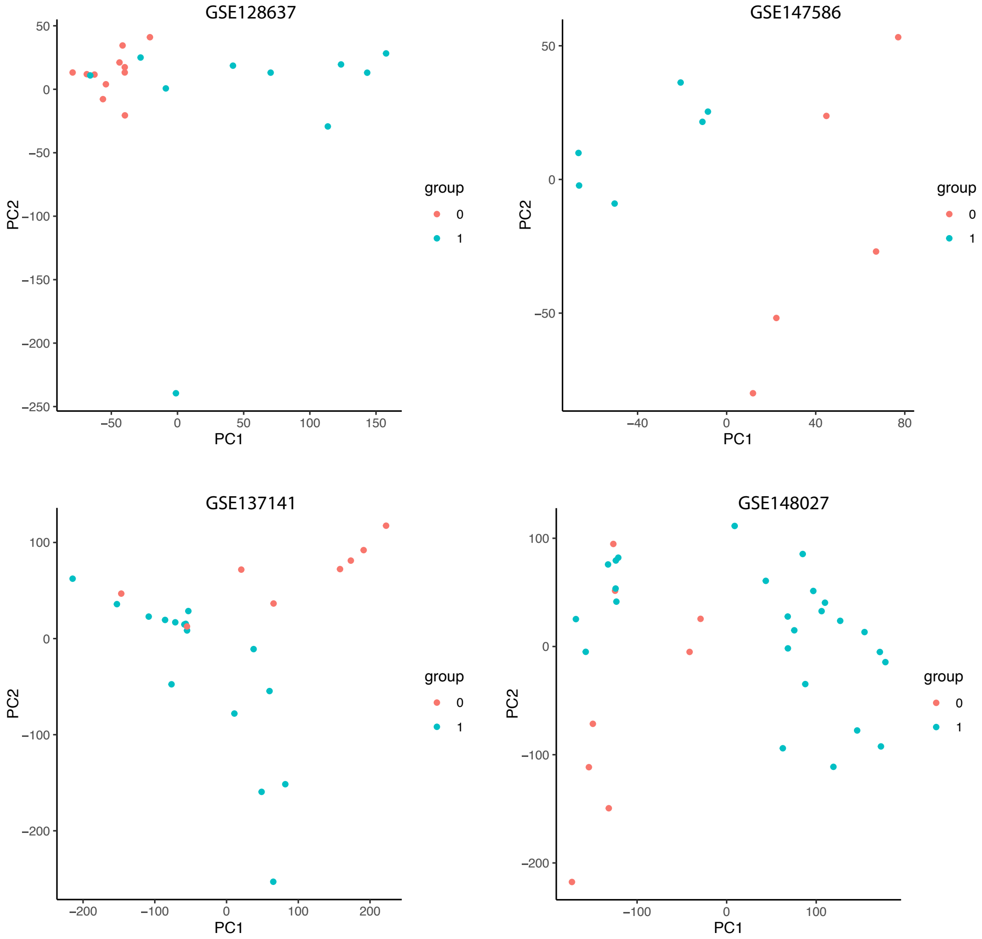

**Figure S2: Meta-analysis scores differentiating HS from control skin.** HS disease gene signature obtained from meta-analysis of skin differentiates healthy skin from HS skin (padj < 0.05). Violin plots of the normalized meta score from MetaIntegrator.

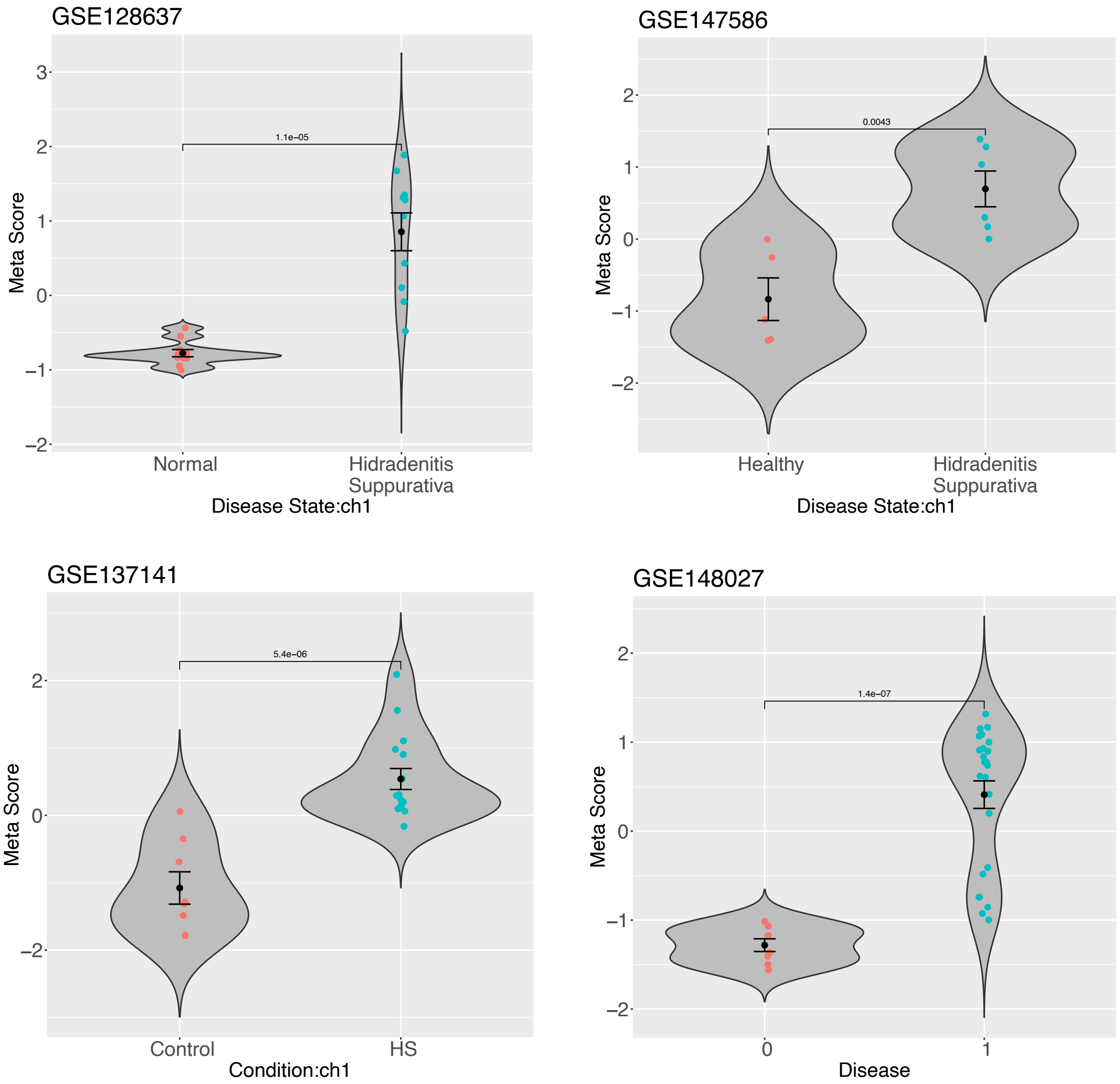

**Figure S3**: **Distribution of drug reversal scores.** Histogram of drug reversal scores of all screened drugs for skin and blood HS disease signatures. Negative reversal scores correspond to drugs predicted to reverse the HS disease signature.

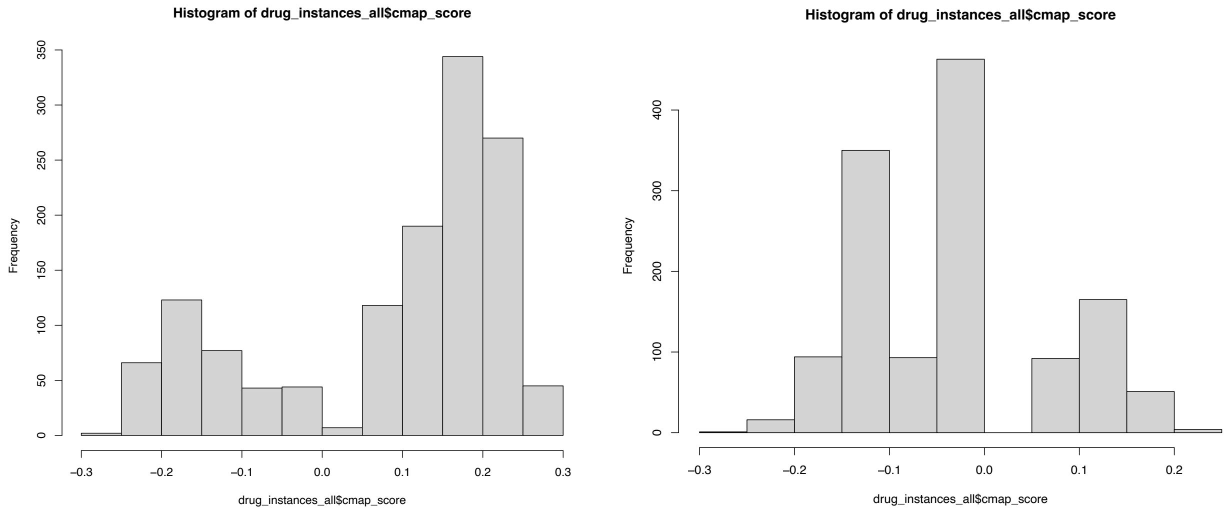

**Figure S4: Aggregate gene-drug reversal signatures for HS skin**

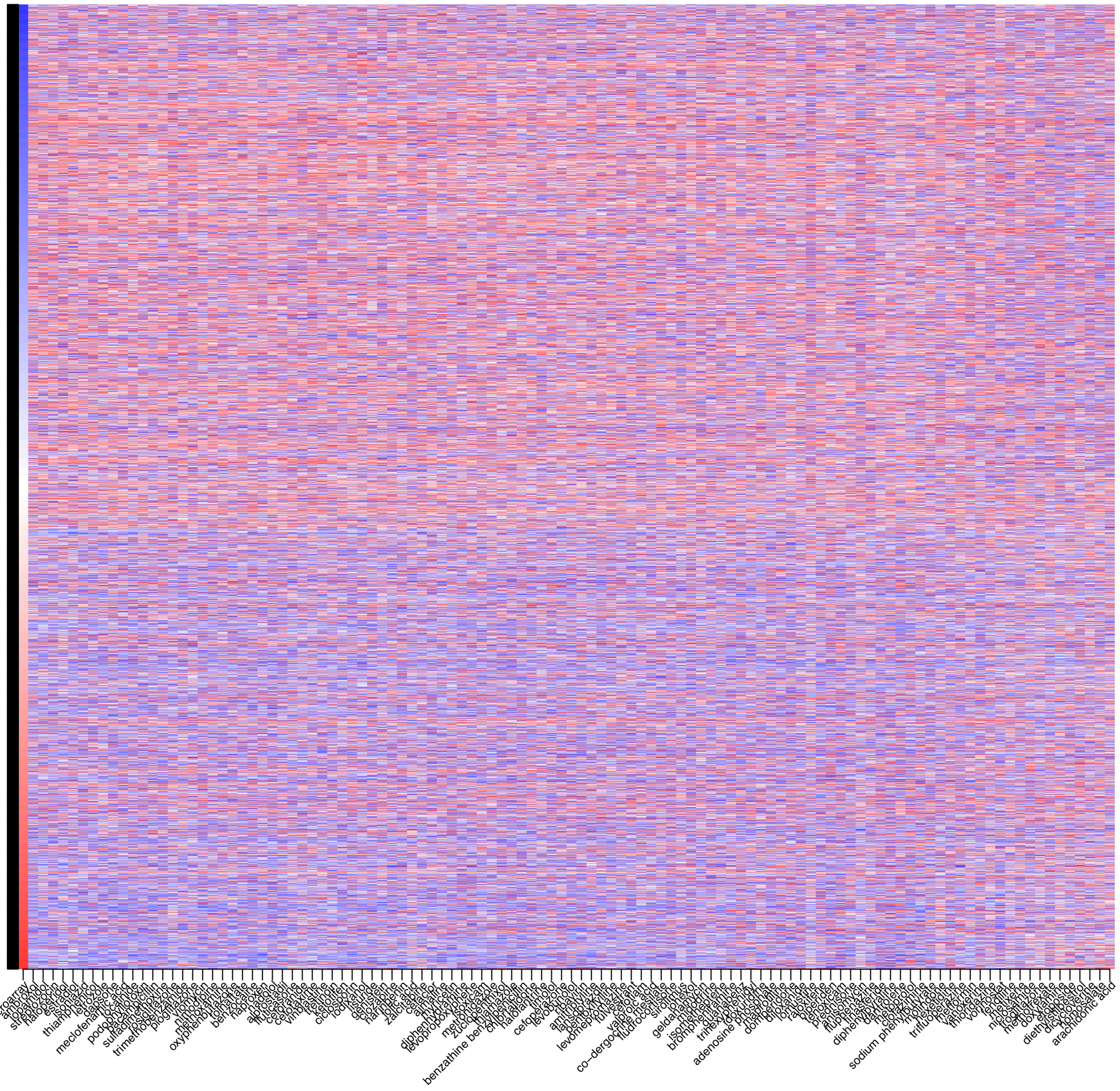

### Figure S5: Aggregate gene-drug reversal signatures for HS blood

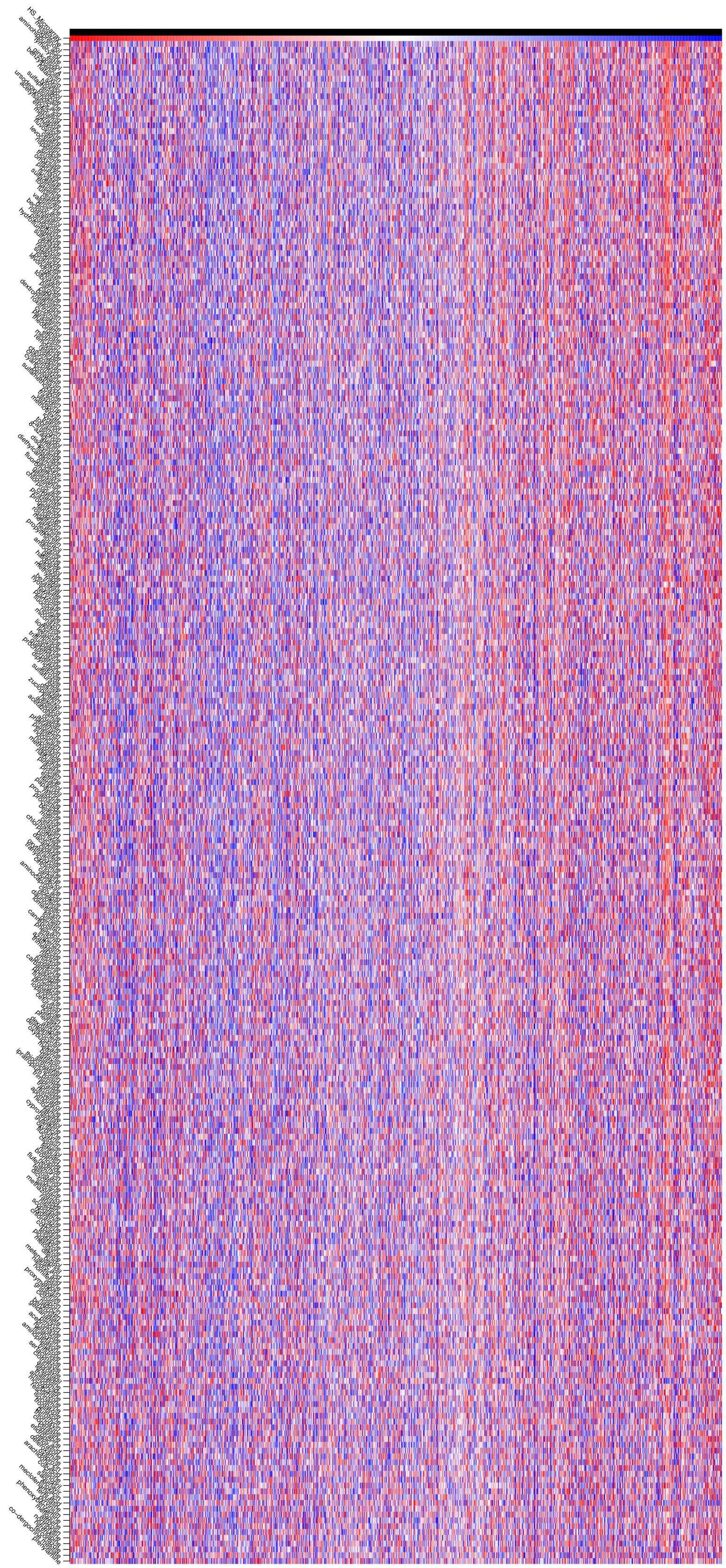

**Figure S6: Drug-protein target interactome for a subset of top predicted drug hits**

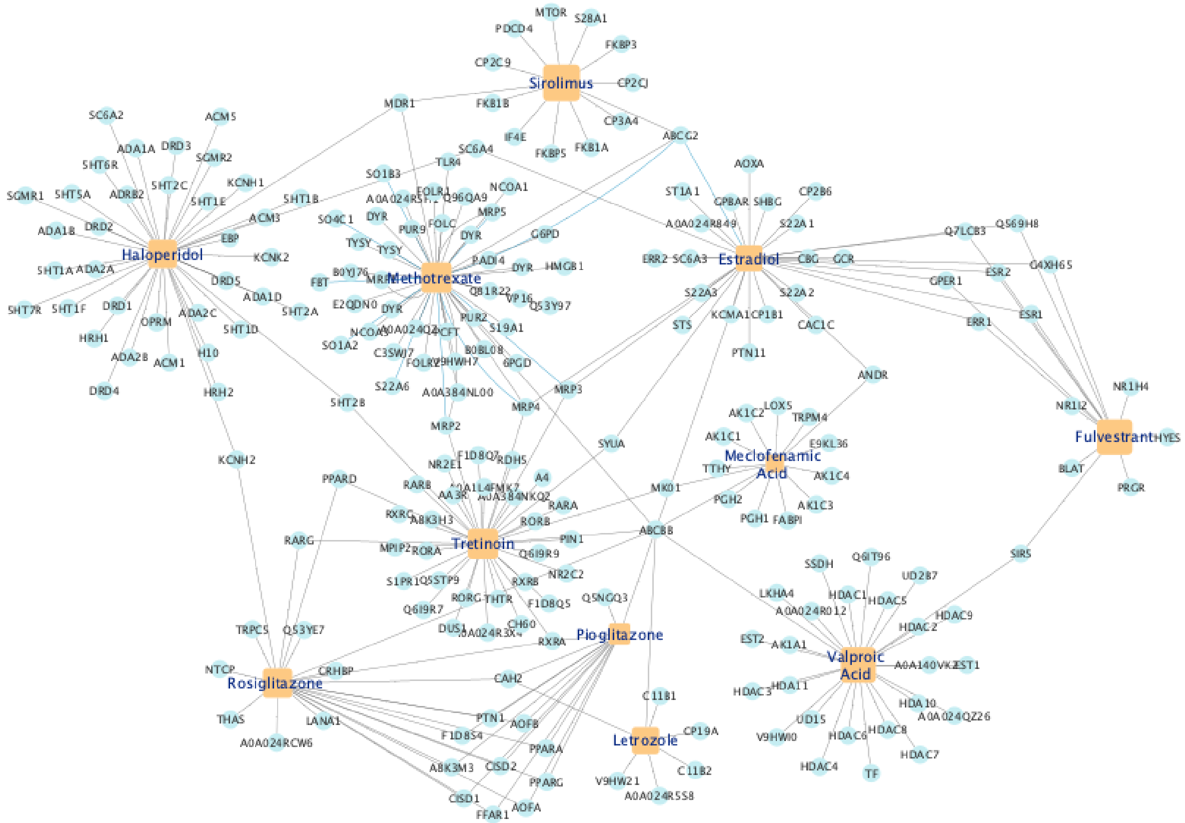

### Figure S7: Cell type abundances from HS skin immune cell scRNAseq.

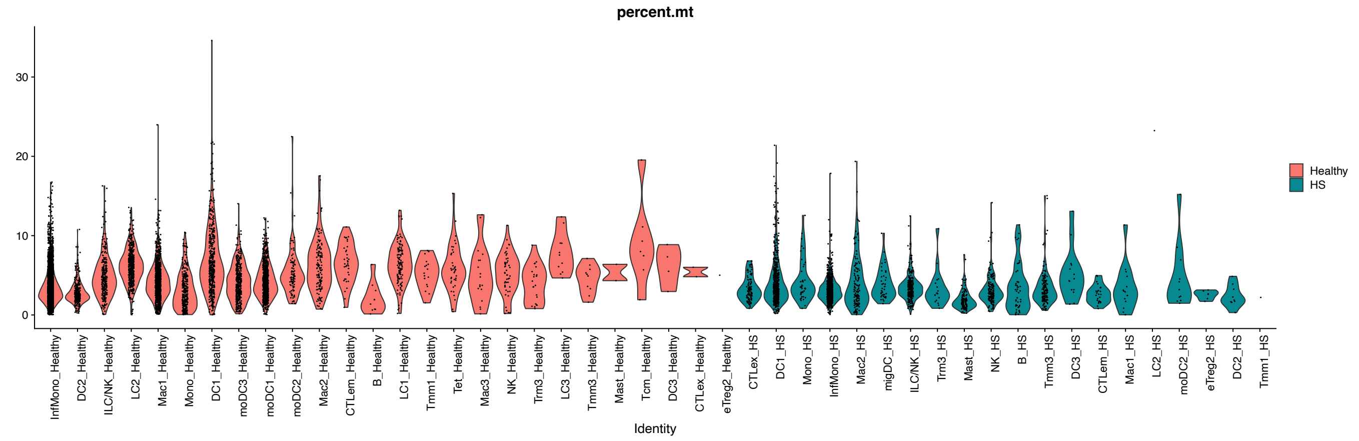
